# The evolutionary traceability of proteins

**DOI:** 10.1101/302109

**Authors:** Arpit Jain, Arndt von Haeseler, Ingo Ebersberger

## Abstract

Orthologs document the evolution of genes and metabolic capacities encoded in extant and ancient genomes. Orthologous genes that are detected across the full diversity of contemporary life allow reconstructing the gene set of LUCA, the last universal common ancestor. These genes presumably represent the functional repertoire common to – and necessary for – all living organisms. Design of artificial life has the potential to test this. Recently, a minimal gene (MG) set for a self-replicating cell was determined experimentally, and a surprisingly high number of genes have unknown functions and are not represented in LUCA. However, as similarity between orthologs decays with time, it becomes insufficient to infer common ancestry, leaving ancient gene set reconstructions incomplete and distorted to an unknown extent. Here we introduce the *evolutionary traceability*, together with the software *protTrace*, that quantifies, for each protein, the evolutionary distance beyond which the sensitivity of the ortholog search becomes limiting. We show that the LUCA set comprises only high-traceable proteins most of which have catalytic functions. We further show that proteins in the MG set lacking orthologs outside bacteria mostly have low traceability, leaving open whether their eukaryotic orthologs have just been overlooked. On the example of REC8, a protein essential for chromosome cohesion, we demonstrate how a traceability-informed adjustment of the search sensitivity identifies hitherto missed orthologs in the fast-evolving microsporidia. Taken together, the evolutionary traceability helps to differentiate between true absence and non-detection of orthologs, and thus improves our understanding about the evolutionary conservation of functional protein networks.

## Introduction

‘How old is a gene?’ is one of the fundamental questions in functional and evolutionary genetics (Capra, et al. 2013). The age of a gene is tightly linked to many of its functional properties. Proteins encoded by old genes tend to evolve slightly slower than younger genes (Wolf, et al. 2009), they are expressed in more tissues (Freilich, et al. 2005), are more central in protein-protein-interaction networks (Kim and Marcotte 2008), and they seem involved in more complex regulatory networks (Warnefors and Eyre-Walker 2011). It, thus, comes as little surprise that gene age is a good proxy for the essentiality of the encoded protein’s function (Gustafson, et al. 2006; Hwang, et al. 2009), and that older genes are more often associated with human diseases (Domazet-Loso and Tautz 2008; Cai, et al. 2009; Maxwell, et al. 2014). Assessing the age of a gene, however, is not trivial (Capra, et al. 2013), as none of the above characteristics can be attributed exclusively to old genes (Wolf, et al. 2009). Instead, age estimates are typically derived from interpreting, for each gene, the phylogenetic distribution of its orthologs (Mirkin, et al. 2003). Under the simplifying assumption that genes are transferred only vertically from ancestor to descendent, the last common ancestor of the two most distantly related species in a phylogeny harbouring an ortholog, approximates the minimal age of the corresponding gene (but see (Doolittle 1999; Gogarten, et al. 2002)). Genes of the same age can then be summarized in phylostrata (Domazet-Loso, et al. 2007), which inform about the lineage-specific evolution of gene repertoires (Ebersberger, et al. 2014), and allow to correlate genetic innovation with major changes during organismal evolution (Slamovits, et al. 2004; Domazet-Loso, et al. 2007; Sestak and Domazet-Loso 2015). The oldest layers in the phylostrata comprise the genes whose orthologs span a considerable or even the full diversity of contemporary life. These genes are likely to hold key position in the metabolic network, and their widespread phylogenetic distribution implies that a loss is detrimental for survival (Mushegian and Koonin 1996). In particular, those genes that can be traced back to the last universal common ancestor (LUCA) (Woese 1998; Goldman, et al. 2013), have been used to deduce the molecular scaffold essential for organismic life (Koonin 2003).

Design of artificial life challenges the evolutionary inferences of a universal genetic repertoire common to – and necessary for – all living organisms (reviewed in (Rancati, et al. 2017)). Only recently, 473 genes from *Mycoplasma mycoides* were determined as the minimal gene (MG) set required, under the most favourable conditions (Koonin 2003), for a self-replicating cell (Hutchison, et al. 2016). Many of these genes have no detectable homologs outside bacteria or even Mycoplasma (Hutchison, et al. 2016) suggesting an evolutionarily recent origin. This is at odds with the expectation that essential genes have a wide phylogenetic spread (Jordan, et al. 2002). Instead, it seems to highlight that also essential gene sets are subject to evolutionary change (Rancati, et al. 2017). For example, a gene responsible for an essential function can be replaced by an unrelated, yet functionally equivalent gene, a process called non-orthologous gene displacement (Koonin, et al. 1996). Alternatively, genes that are essential in one organism may not be essential in another (Liao and Zhang 2008; Koo, et al. 2017), e.g. because its metabolic network has become more robust by evolving redundancy, or because the metabolic network was rewired to bypass essentiality of individual proteins (Kim, et al. 2010; Rancati, et al. 2017). Taken together, this implies that the *Mycoplasma mycoides* MG set represents only a minor step towards unravelling the universal building plan of organismic life.

However, sequence similarity used to identify orthologs in present day gene sets decays with time (Dayhoff 1978). Ultimately, a twilight zone (Doolittle 1981) is hit where two related proteins are no longer similar enough to infer common ancestry (Dayhoff 1978; Rost 1999).

The time to reach the twilight-zone varies between proteins and depends on their sequence composition as well as their substitution rate (Dayhoff 1978), but not on their essentiality (Hurst and Smith 1999; Hirsh and Fraser 2001). This links the accuracy of the gene age assessment to the sensitivity of the ortholog prediction methods, where a low sensitivity will obtain incomplete phylogenetic profiles. As a consequence, the sharing of essential genes between distantly related or fast evolving species will be overlooked, and gene ages will be underestimated (Luz, et al. 2006; Moyers and Zhang 2015, 2016, 2017). The risk of misinterpreting the evolutionary past is therefore high (Liebeskind, et al. 2016; Martin-Duran, et al. 2017). While individual approaches exit that aim at delineating the evolutionary distance beyond which orthologs no longer share a significant sequence similarity (e.g. (Moyers and Zhang 2016)), standardized solutions cast into a dedicated software are not yet at hand. Our understanding to what extent the same functions in contemporary species are conveyed by the same evolutionarily old genes is incomplete to an unknown extent.

Here we introduce for each protein its (*evolutionary) traceability* informing over what evolutionary distances sequence similarity should suffice for an ortholog identification. Using the yeast as a showcase, we find that genes with high traceabilities are enriched for catalytic functions in the cell metabolism. The subset of yeast genes whose evolutionary origins have been dated back to LUCA are almost all of high traceability. A substantial fraction of the yeast genes, among them many with essential functions, however, have low traceabilities. This is a clear sign that the sensitivity of an ortholog search does not suffice to identify orthologs in distantly related species. These findings are further corroborated by a traceability analysis of the MG set. The vast majority of the MG-set proteins that appear confined to bacteria show low traceabilities indicating a high chance that orthologs in more distantly related species have been overlooked. Finally, we demonstrate exemplarily for yeast Rec8, a protein essential for recombination, how a traceability-informed increase of the ortholog search sensitivity can lead to the identification of hitherto overlooked representatives of Rec8 in the fast-evolving microsporidia.

## New Approaches

### protTrace: A simulation-based workflow to estimate evolutionary traceability of a protein

protTrace is a simulation-based framework that determines for a user-defined protein, the seed-protein, its traceability as a function of evolutionary time. The procedure comprises four main steps – (1) Parameterization of a site-specific evolutionary model, (2) simulation of protein sequence evolution, (3) the calculation of the traceability, and optionally (4) the display of the traceabilities on a reference tree. The detailed workflow is represented in fig. 1 and in supplementary fig. S1A.

**Figure 1.**
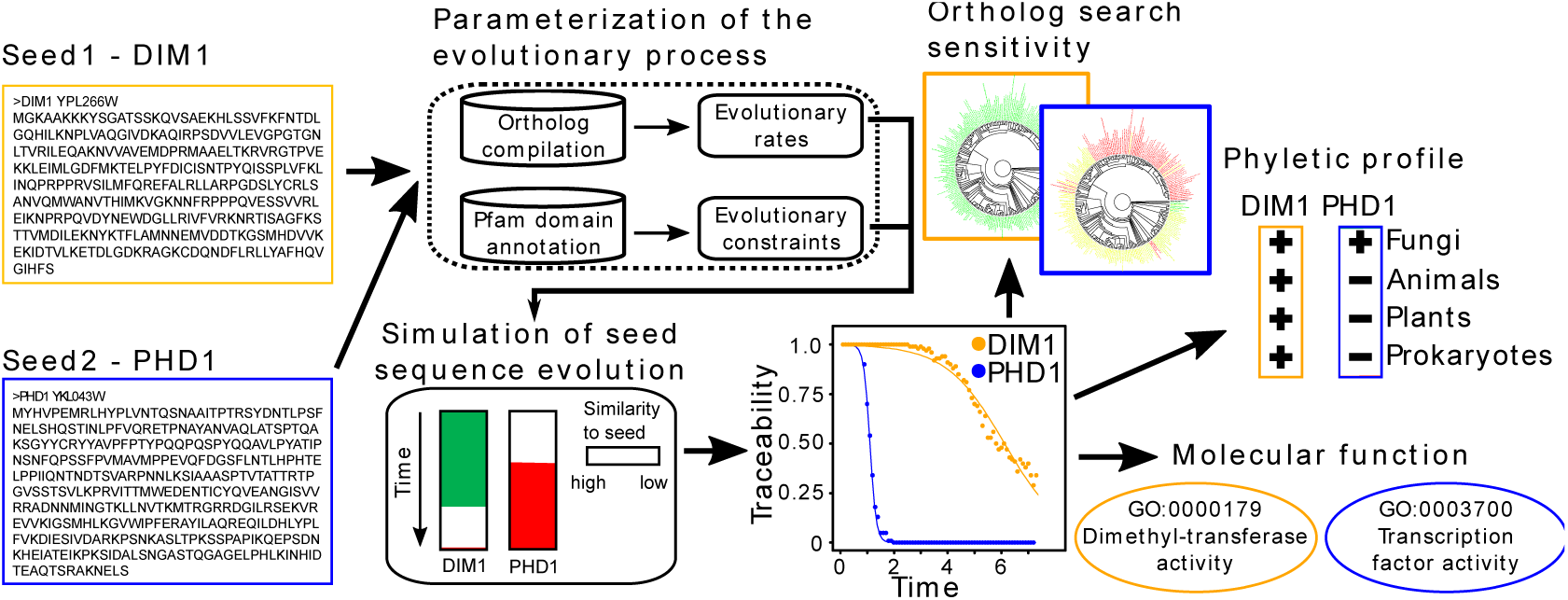
Workflow to assess the evolutionary traceability of a protein. The framework is exemplified for yeast PHD 1(blue) and DIM1 (yellow). For each seed-protein, we use a simulation-based approach to infer its traceability, *Ti(t)*, that is defined on the interval [0,1]. From its traceability graph and the evolutionary distance to any target species, the traceability of the seed in the target species can be extracted. Relating this information to (i) a species tree highlights taxa where the ortholog search sensitivity becomes limiting (red clades), (ii) phylogenetic profiles identifies cases where orthologs might have been overlooked, and (iii) the gene ontology identifies molecular functions that coincide with low traceability.

### Step 1 - Parameterization of the evolutionary process

First, ProtTrace infers the evolutionary characteristics of the seed-protein. We compile a group of orthologs, *O_seed_* for the seed-protein. ProtTrace facilitates the use of pre-compiled orthologs from OMA (Altenhoff, et al. 2015), InParanoid (Ostlund, et al. 2010) and from orthoDB (Zdobnov, et al. 2017). Optionally, a targeted ortholog search with HaMStR (Ebersberger, et al. 2009) can be employed. In the next step, the orthologous sequences are aligned with MAFFT v7.304 (Katoh and Toh 2008), and a maximum likelihood tree, *T_seed_*, is computed with RAxML v8 (Stamatakis 2014). The resulting tree and the multiple sequence alignment (MSA) are then used to determine the evolutionary parameters of the proteins. A maximum parsimony algorithm infers the seed protein-specific insertion and deletion (indel) rates. The procedure is illustrated in supplementary fig. S1B, and Supplementary fig. S2A shows the distribution of the insertion rates exemplarily for the yeast protein set. Finally, the insertion and deletion lengths of one most parsimonious solution are used for estimating *p*, the parameter of the geometric indel length distribution.

In a phylogenomic setting the evolutionary parameters are inferred for many seed-proteins, e.g. all proteins encoded in a species’ genome. To account for different absolute substitution rates between the individual seed-proteins, we introduce a rate scaling factor *κ_seed_* (Eq. 1). We compute *κ_seed_* for each seed-protein as

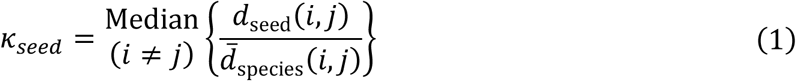

where *d_seed_(i*,*j)* is the maximum likelihood distance, inferred from *T_seed_* for species *i* and *j*, and *d_Species_*(*i*,*j*) is the average maximum likelihood distance across all pair-wise OMA orthologs for the two species *i* and *j*. *κ_seed_* is then the median of the ratio inferred from all species pairs *i*, *j* in *O_seed_*. Supplementary fig. S2B shows the distribution of *κ_seed_* exemplarily for the yeast protein set.

With *hmmscan* (Finn, et al. 2015) (parameters: ‐‐*notextw* and -*E 0.01)* we identify regions in the seed protein representing Pfam-A (Finn, et al. 2016) domains. From the corresponding profile hidden Markov models of the Pfam domains we extract the information for a site-specific domain constraint on the evolutionary process (Koestler, et al. 2012).

### Steps 2 – 3 - Simulation of protein sequence evolution and calculation of the traceability curve

Once the evolutionary model is fully parameterized, ProtTrace uses REvolver (Koestler, et al. 2012) to simulate the evolution of the seed protein in time steps of 0.1 substitutions per site. After each step, the simulated sequence serves as a query for a BLASTP (Altschul, et al. 1997) search against the full protein set of the species the seed-protein was derived from (seed species). If the seed-protein sequence is identified as one of the top five hits, the success is marked with a ‘1’, otherwise a ‘0’ is noted. Repeating the simulation 100 times yields for each time step *t_i_* a fraction of successes that approximates *Ti(t_i_)*, the traceability index of the seed-protein as a function of evolutionary time *t*. We fit the inverse of a non-linear least square logistic growth curve to this data (Eq. 2) using the non-linear least square (*nls*) package in R.

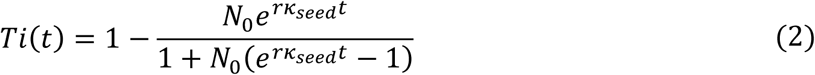

We estimate the parameters *N_0_*, and *r*, the rate change of traceability, from the data.

### Step 4 - Tree display

*Ti(t)* describes the traceability index of the seed protein as a function of time. To provide an intuitive overview for each seed protein, in which species the sensitivity of an ortholog search could become an issue, protTrace can display the traceability information along a species phylogeny (*see* supplementary fig. S3). Here, the colour of the leaf labels indicates the traceability of the seed-protein in the respective species.

### Code availability

ProtTrace is available from https://github.com/BIONF/protTrace.git.

## Results and Discussion

### The evolutionary traceability of the yeast gene set

Yeast (*Saccharomyces cerevisiae*), as a genetically and functionally well characterized model organism, provides an excellent starting point for exemplifying concept and implications of protein traceability (fig. 1). We determined the traceability index (*Ti(t)*) for 6,352 yeast proteins in 232 target species representing all three domains of life (*see* supplementary tables S1 and S2). If an ortholog group for a yeast protein comprised less than four sequences, we used as default the mean of the indel rate distribution across the entire protein set (0.08) (*see* supplementary figure S2A), and set the parameter *p* of the indel length distribution to 0.25. If no ortholog was detected for a seed-protein, we used the mean of the scaling factor distribution across all yeast proteins (*κ_mean_*=1.57) as the default value (*see* supplementary figure S2B). The distribution of the resulting traceabilities is shown in Supplementary figure S4. Orthologous groups based on OMA and complemented with HaMStR (see Methods), or compiled with OrthoDB obtained highly correlated results (r = 0.92; data not shown). The choice of the ortholog search method has therefore almost no impact on the traceability estimate, and we used the traceabilities obtained from the OMA / HaMStR approach for the remainder of the analysis. Likewise, there was virtually no impact on the traceability estimates if we recruited the orthologs for estimating the evolutionary parameters from species across the entire tree of life or only from fungal species (*see* Supplementary fig. S5). Fig. 2A displays the traceabilities of the yeast proteins exemplarily for four eukaryotes, one archaeon and one bacterium. For 2,040 proteins, the traceabilities decreases only very slowly with increasing evolutionary distance between yeast and the target species (*Ti(t)≥0.95* for all target species). As we cover the full phylogenetic diversity in the tree of life, rate and pattern of evolutionary sequence change for these proteins should not hinder ortholog detection in any extant species. For the remaining 4,312 sequences, phylogenetic distance and the evolutionary rate of the target species jointly determine protein traceability. When moving from the closely related fungus, *A. gossypii*, to archaea and bacteria, the number of proteins with a traceability of 0 increases by an order of magnitude (fig. 2A). Likewise, traceabilities are substantially smaller in the microsporidium *E. cuniculi*, an obligate intracellular parasite closely related to fungi (Thomarat, et al. 2004), than in human and Arabidopsis that belong to different kingdoms. This is an effect of the extraordinarily high substitution rate in the microsporidian lineage, which is among the highest across all eukaryotes (Slamovits, et al. 2004).

**Figure 2.**
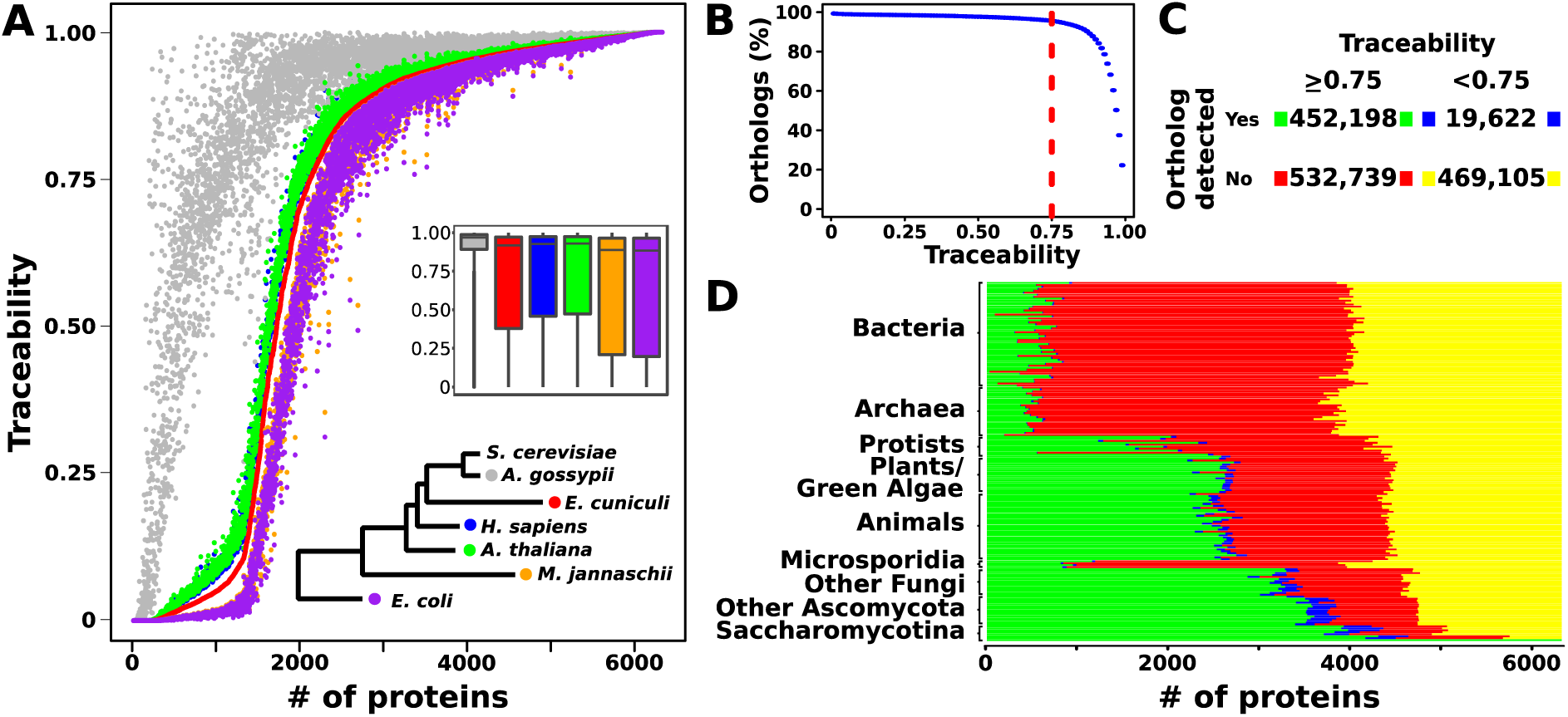
The evolutionary traceability of yeast proteins. **A**, Traceabilities for 6,352 yeast proteins in *Ashbya gossypii* (Fungi), *Encephalitozoon cuniculi* (Microsporidia), *Homo sapiens* (Metazoa), *Arabidopsis thaliana* (Viridiplantae), *Methanocaldococcus jannaschii* (Archaea) and *Escherichia coli* (Bacteria) individually, and as a box plot. Proteins are ordered according to their traceability in *E. cuniculi.* The color code is specified in the phylogenetic tree. **B**, Cumulative distribution of the detected yeast orthologs relative to protein traceability. 95% of the detected orthologs coincide with A traceability of 0.75 or above (red line). **C**, Relation between results of the ortholog search and protein traceability. **D**, shows the per-species results with the colour code following C.

We next calibrated the traceability index such that it informs in real data about the evolutionary distance beyond which orthologs are too diverged to be detected with BlastP based ortholog predictors. We searched for orthologs in the 232 target species using each of the 6,352 proteins as a query and tabulated the number of query-species pairs in which at least one ortholog was found. In 95% of the cases where an ortholog was detected, the traceability was at least 0.75 (Fig. 2B). This observation leads us to hypothesize that, when the traceability is below 0.75, an ortholog search is bound to fail. If an ortholog exists, it has likely diverged beyond recognition. On the basis of this hypothesis, we distinguish two scenarios for the cases where no ortholog was identified (Fig. 2C). For the 53% of the cases where the traceability is 0.75 or above, we conclude that the ortholog is genuinely absent, as we should be able to detect it otherwise. For the remaining 47%, the traceabilities do not reach the threshold of 0.75, and these cases occur in almost all target species (Fig. 2D). In other words, in almost half of the cases where we do not find an ortholog for a yeast protein, we cannot distinguish between true absence or an insufficient search sensitivity.

We are aware of one previous study that performed an in-silico evolution of the yeast protein set (Moyers and Zhang 2016). In this study, the authors inferred their constraints on the evolutionary process for each yeast protein from the alignment of orthologs of five sensu stricto yeast species. Unfortunately, Moyers and Zhang (2016) did not link their findings to the actual phylogenetic profiles of the yeast proteins, making a comparison to our study hard. We therefore reproduced their analysis in part. Moyers and Zhang (2016) used site specific substitution rate scaling factors inferred with TreePuzzle (Schmidt and von Haeseler 2007) as information to constrain the evolutionary process in a site-specific manner. We recreated these constraint vectors, once with the original approach by Moyers and Zhang (2016) using the five sensu stricto yeast sequences, and once with an alignment using orthologs selected from the full diversity of fungi. This revealed that the phylogenetic diversity of the input alignment has a strong effect on the constraint pattern. When using the sensu stricto yeast orthologs, on average 80 *%* of the alignment sites are assigned a relative rate of 0. Such positions remain unchanged during evolution. In contrast, when using the phylogenetically diverse training data, on average only about 15% of the alignment sites get assigned a relative rate of 0 (*see* supplementary figure S6). Thus, the evolutionary constraint information –and as a consequence the simulated change of the protein over time— changes with the underlying training data. This raises the question about the optimal collection of training data. In the particular case of the simulated yeast protein evolution (Moyers and Zhang 2016), it appears that the use of the closely related yeast sequences for inferring the site-specific rate scaling factors puts a too harsh constraint on the evolutionary process (*see* supplementary figure S6). Using our terminology, this is bound to result in an overestimated traceability, an aspect that the authors have noted themselves (Moyers and Zhang 2017).

### Unobserved domain constraints result in underestimated traceabilities

The integration of traceability and ortholog search for the yeast proteins reveales that we sometimes (5%) detect an ortholog although the traceability of the seed-protein predicts that we should not. Reducing the traceability cut-off has little effect on this number (fig. 2B). The underlying causes that can explain such a discrepancy between the traceability estimate and the outcome of an ortholog search are diverse (*see* supplementary text). On the one hand, overestimates of the protein-specific evolutionary rates can artificially decrease the traceabilities – although protTrace is considerably robust with respect to variation in the rate estimates (*see* supplementary fig. S7). On the other hand, spurious ortholog assignments can mimic the presence of an ortholog where there is none, an artefact that is obviously hard to control. One main factor determining a protein’s traceability, however, is its Pfam domain content (Finn, et al. 2016), as protTrace exploits the characteristic sequence features of Pfam domains to deduce functional constraints on the evolutionary process (Koestler, et al. 2012). In the yeast data, 1,255 out of 6,352 proteins do not harbour any Pfam domain. In the simulations that underlie the traceability estimates, these proteins evolve without position-specific constraint, and correspondingly have low traceabilities (*see* supplementary fig. S8). This implies that protTrace, if information concerning local constraints on the sequence-specific evolutionary process is not provided, can underestimate the traceability of a protein. Fig. 3 describes an illustrative example. The yeast protein MRS2 is a mitochondrial inner membrane Mg^2+^ transporter (Wiesenberger, et al. 1992), and its traceability outside fungi is substantially below the critical value of 0.75 (*see* supplementary table S2). The low traceability estimate coincides with the absence of any Pfam domain in the MRS2 sequence (fig. 3A). However, we find yeast MRS2 orthologs across the entire eukaryotic domain (fig. 3B), suggesting that the traceability estimate by protTrace is too low. A multiple sequence alignment of these orthologs resolves the issue (fig. 3C). MRS2 harbours evolutionarily highly conserved domains, which have no resemblance in Pfam, and thus could not be taken into account during the traceability estimation. Notably, when we generate a custom pHMM from the MRS2 alignment and use this as a constraint model for the sequence simulation within protTrace, the mean traceability of this protein increases from 0.07 to 0.97 (data not shown).

**Figure 3.**
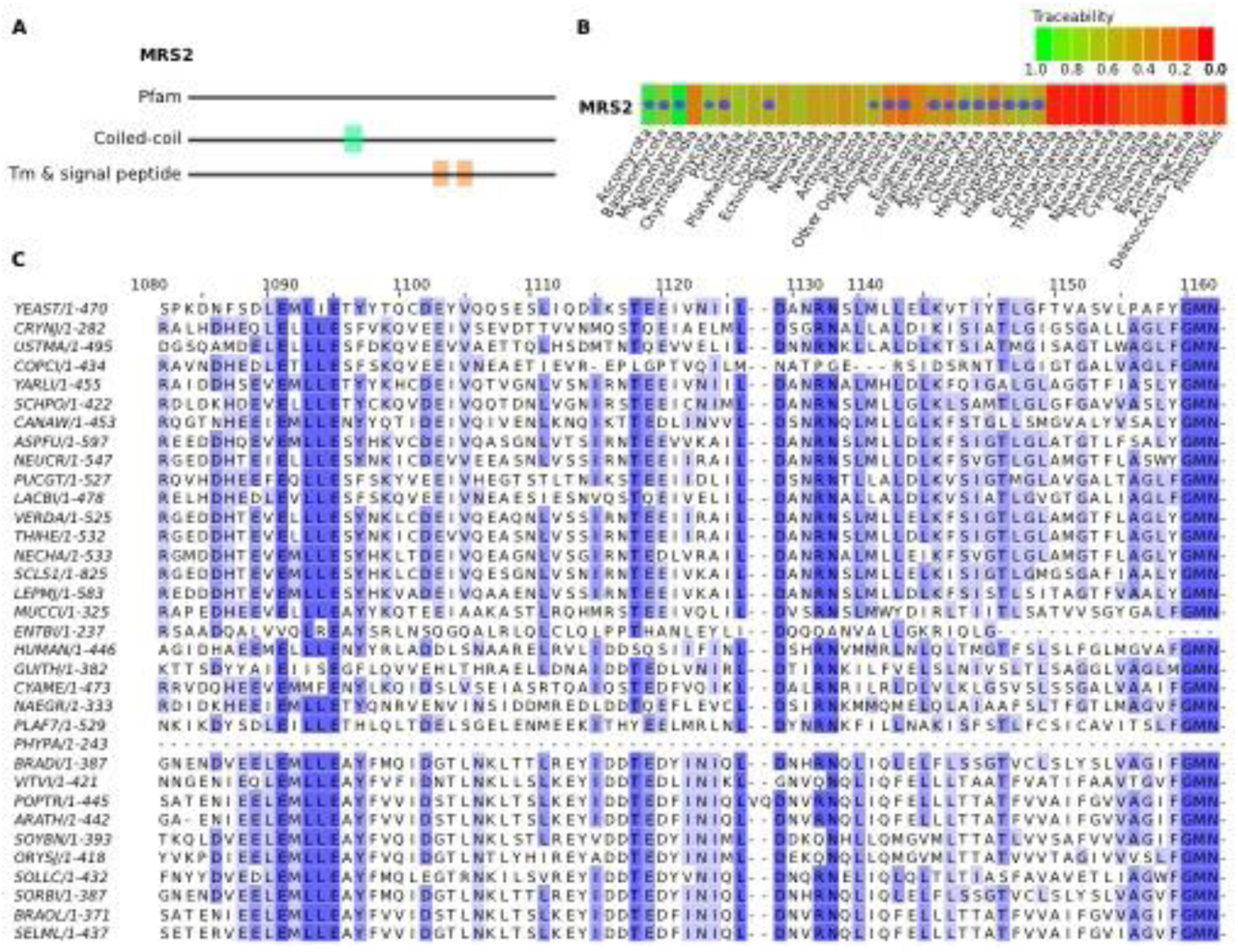
Missing information about domain constraints result in underestimated traceabilities: The yeast mitochondrial inner membrane Mg2+ transporter MRS2. **A**, MRS2 displays no significant hit against any Pfam domain and contains as sole features a central coiled-coil domain and two transmembrane domains. **B**, The phylogenetic profile of MRS2 reveals the existence of orthologs across the entire eukaryotic kingdoms despite a predicted low traceability. The presence of an ortholog in a given species is indicated by a dot. The cell colour represents protein traceability. **C**, Section of the MRS2 alignment considering orthologs from different representatives across the eukaryotic tree of life. The selected region shows exemplarily for the entire alignment that MRS2 orthologs share conserved sequence motifs that most likely are associated with the functionality of this protein as a Mg2+ membrane transporter. As these conserved domains are not represented in a Pfam domain, protTrace cannot consider the corresponding evolutionary constraints during its simulation.

We expect that there are other cases like MRS2. Already in the past three years, and from Pfam release 27 (Finn, et al. 2014) to Pfam release 29 (Finn, et al. 2016), has the number of models increased from 14,831 to 16,295, and it is likely that this catalogue is still not complete. Thus, a fraction of yeast sequences might evolve under a hitherto undescribed evolutionary constraint, and protTrace, when using only Pfam constraints, will underestimate its traceability. In such cases it is advisable to start protTrace with the option to extract site-specific constraints on the evolutionary process directly from a multiple sequence alignment of ortholgos, similar to previous approaches (Alba and Castresana 2007; Moyers and Zhang 2015, 2016). It might be interesting to note that discrepancies between traceability and evolutionary profile, as exemplified by MRS2, can be easily applied to automatically screen for further such instances, where a functional domain is currently not described in Pfam.

### Traceability and sub-cellular localization are linked

Protein traceability informs whether or not the sensitivity of an ortholog search is sufficient to accurately determine the phylogenetic profile of a protein even in distantly related species. ofInitial evidence that this measure can provide an alternative view on the interpretation of conservation patterns of orthologs across species comes from the analysis of proteins with different sub-cellular localization. It was recently reported that membrane proteins, and even more so extracellular proteins, have sparser phylogenetic profiles and fewer detected orthologs than intracellular proteins (Sojo, et al. 2016). Both was taken as evidence for a rapid evolutionary turnover of membrane and extracellular proteins. We addressed the same issue from a view point of protein traceability. We classified the yeast proteins into three groups - membrane proteins, extracellular proteins and intracellular proteins – as described previously (Sojo, et al. 2016). For every protein, we computed the mean traceability across the 232 target species and found that extracellular proteins have overall lower traceabilities compared to the membrane-bound and intracellular proteins (fig. 4). For the latter two groups the differences in the mean traceabilities were less pronounced. Yet, the distribution for the membrane proteins displays a higher fraction of proteins with mean traceabilities below 0.5. In the light of these results we expect that an ortholog search more often misses a distantly related ortholog for extracellular and membrane proteins than for intracellular proteins. This is perfectly in line with the differences in their observed evolutionary conservation of the three groups as reported by Sojo et al (2016). We therefore conclude that the particular evolutionary behaviour of proteins with different sub-cellular localizations, and their differences in the traceability together account for the observed differences in their evolutionary conservations.

**Figure 4.**
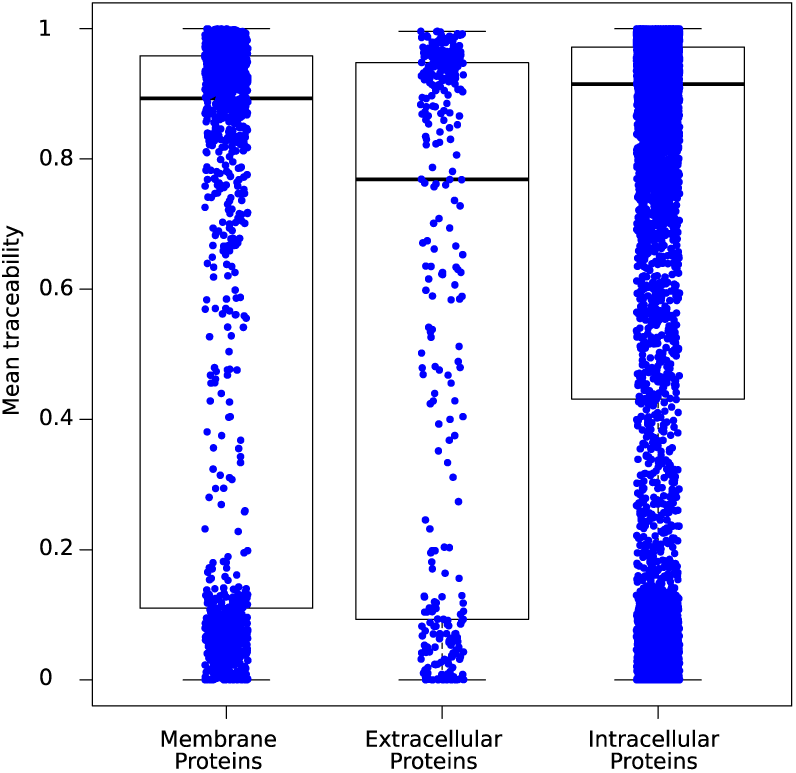
Boxplot of the mean traceabilities across 232 species for the yeast proteins in dependence of their subcellular localization. The individual data points underlying the boxplot are represented in blue.

### Protein traceability, molecular function, and gene age estimates are linked

Earlier studies have reported the rapid evolution of proteins that are part of the immune defense, reproductive processes, cell adhesion and transmembrane transport (Swanson and Vacquier 2002; Panhuis, et al. 2006; Voolstra, et al. 2011). Exemplarily for yeast, we evaluated the link between the traceability of a protein and its function, as represented in the assignment of Gene Ontology (GO) terms (Ashburner, et al. 2000). We split the 6,352 yeast proteins into three bins based on their predicted traceabilities in *E. coli* ([0.75 - 1]: 3947 proteins; [0.25 - 0.75]: 742 proteins; [0 - 0.25]: 1663 proteins). A subsequent characterization with Gorilla (Eden, et al. 2007; Eden, et al. 2009) and visualization of the results with Revigo (Supek, et al. 2011) reveals that GO terms are not identically distributed across the three categories (*see* supplementary fig. S9). The 3,947 high-traceability yeast proteins *(Ti(E.coli)≥0.75)* are significantly enriched for catalytic functions (*see* supplementary fig. S9A). Among these, we find 98% of the 980 yeast enzymes annotated by the Enzyme Commission (EC). Regulatory functions, in turn, are overrepresented in the group of 742 proteins with intermediate traceabilities between 0.75 and 0.25 (*see* supplementary fig. S9B). The proteins with traceability below 0.25 are preferentially involved in cell aggregation and cell reproduction (*see* supplementary fig. S9C). Altogether, we find that 17% essential proteins (Giaever, et al. 2002) and 70% of the yeast transcription factors have traceability below 0.75 (*see* supplementary table S3). The low traceability implies that the orthology between regulatory proteins, but also between proteins of other essential functionalities are difficult to detect across distantly related species. Consequently, such functions should be underrepresented in the reconstructions of ancient gene sets, not because they are necessarily evolutionary younger, but because information about their evolutionary ancestry decays rapidly.

The 1,203 yeast proteins that are represented in the reconstructed gene set of LUCA (Goldman, et al. 2013) exactly match this prediction. They are almost exclusively (96%) recruited from the high-traceability bin. They comprise about half (47%) of all EC annotated yeast enzymes, but merely 4% of the 245 transcription factors with a known binding site (de Boer and Hughes 2012). When taken as face value, this observation translates into a complex evolutionary scenario: The molecular ‘hardware’ of contemporary species, consisting mainly of enzymes, was largely established first already in LUCA. The regulatory ‘software’, however, was either independently re-built, or invented multiple times on individual evolutionary lineages (Charoensawan, et al. 2010). In the light of the limited traceability of proteins involved in regulation, it is worth considering a second, more parsimonious explanation. In addition to enzymatic activity, other essential functions might have had a unique genesis early in organismal evolution. However, because rate and pattern of evolutionary sequence change for some of these proteins has eradicated all traces of their ancient origins, it appears as multiple independent inventions of the same function on individual evolutionary lineages.

### Evolutionary traceability of the bacterial minimal gene set Syn3.0

A reanalysis of the data generated by the Artificial Life Project (Hutchison, et al. 2016) corroborates the findings from the previous section. The artificial life project synthesized a self-replicating bacterium (Syn3.0) on the basis of only 438 protein-coding genes from the bacterium *Mycoplasma mycoides* (Hutchison, et al. 2016) (MG set). This collection of essential genes comes close to what Koonin (2003) referred to as an absolute minimal gene set, i.e. the set of genes that an organism requires under the most optimal conditions. One could naively assume that many of these genes are essential for cellular life in general, and are thus conserved across the tree of life. As a consequence, they should be represented in the gene set assigned to LUCA. To assess the phylogenetic distribution of the 438 genes, we replaced the unidirectional Blast search performed by Hutchison, et al. (2016), that does not inform about the precise evolutionary relationships of the identified homologs, with an ortholog search (fig. 5 and see supplementary table S4). This revealed that 170 of these genes have no detectable ortholog outside *Mycoplasma*, and for 149 genes the exact biological function is unclear. On the first sight this might imply that *Mycoplasma* has evolved its own path to organismal functionality reflecting that a set of genes essential for one species must not be essential for another organism (Gerdes, et al. 2003; Koo, et al. 2017). However, we found that 60 proteins in the MG set have traceabilities below 0.75 in any tested species outside *Mycoplasma.* Among these are the majority of proteins with unknown functions (41/65), and additionally 15 of the 84 proteins with only a generic function assigned (fig. 5). Whatever essential tasks these 60 proteins have, it may be premature to mark them as *Mycoplasma*-specific inventions. Instead, we hypothesize that their low traceability blurs the evolutionary link to related proteins with the same function in other organisms. Given their participation in fundamental cellular functioning, it is tempting to speculate that these proteins can provide relevant hints towards the nature of the ‘software’ that appears missing in the current reconstructions of the LUCA gene set.

**Figure 5.**
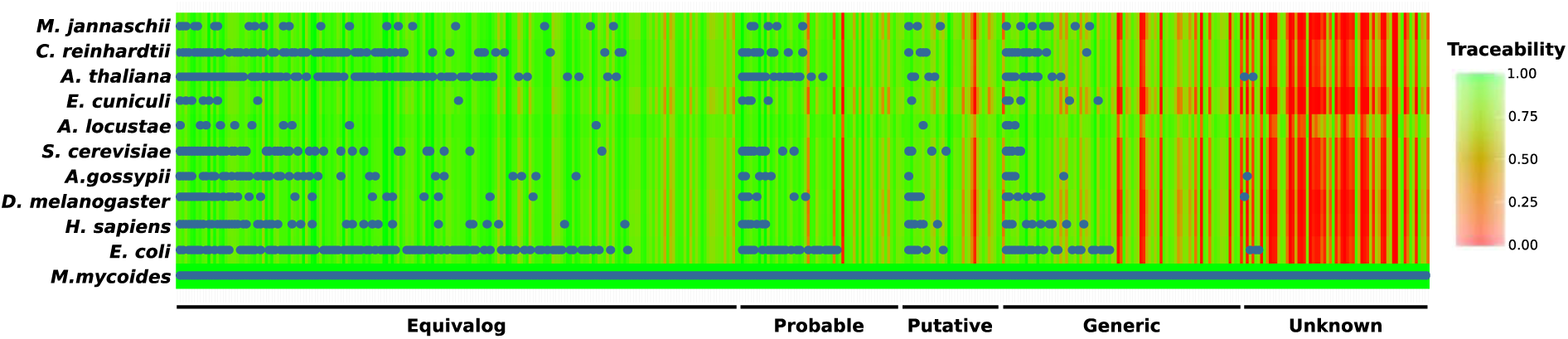
Phylogenetic distribution and traceability profile for the Syn3.0 minimal gene set. The background colour gives the information of protein traceability ranging from green (high traceability) to red (low traceability). The categorization according to the functional annotation status of the individual proteins was adapted from (Hutchison, et al. 2016).

### Protein traceability limits ortholog identification in the fast-evolving microsporidia

Microsporidia, intracellular parasites closely related to fungi (Corradi and Keeling 2009) are a hallmark example that a low traceability can indeed result in essential genes being overlooked. All microsporidia analysed so far share two characteristics: First, their genomes harbour between 2,000 and 4,000 genes, due to an ancient radical reduction in genome size (Slamovits, et al. 2004), and second their proteins evolve extraordinary fast. While the first characteristic makes it tempting to generally equate a non-detection of an ortholog to a yeast protein with a gene loss, the high evolutionary rate of microsporidia indicates that a low traceability may be another reason for the lack of orthologs. Katinka et al (2001) and Cuomo et al. (2012) showed that key metabolic functions, e.g. the fof1-ATPase complex, fatty acid synthesis, the tricarboxylic acid (TCA) cycle and the formation of peroxisomes are absent in microsporidia (Katinka, et al. 2001; Cuomo, et al. 2012). We determined the phylogenetic profiles for the corresponding yeast proteins and could confirm that for many proteins no ortholog was detectable in our microsporidian representatives (fig. 6A and *see* supplementary table S5). For most of these proteins, the traceabilities in microsporidia are in the range of 0.9 and above. This indicates that the corresponding genes have indeed been lost on the microsporidian lineage.

**Figure 6.**
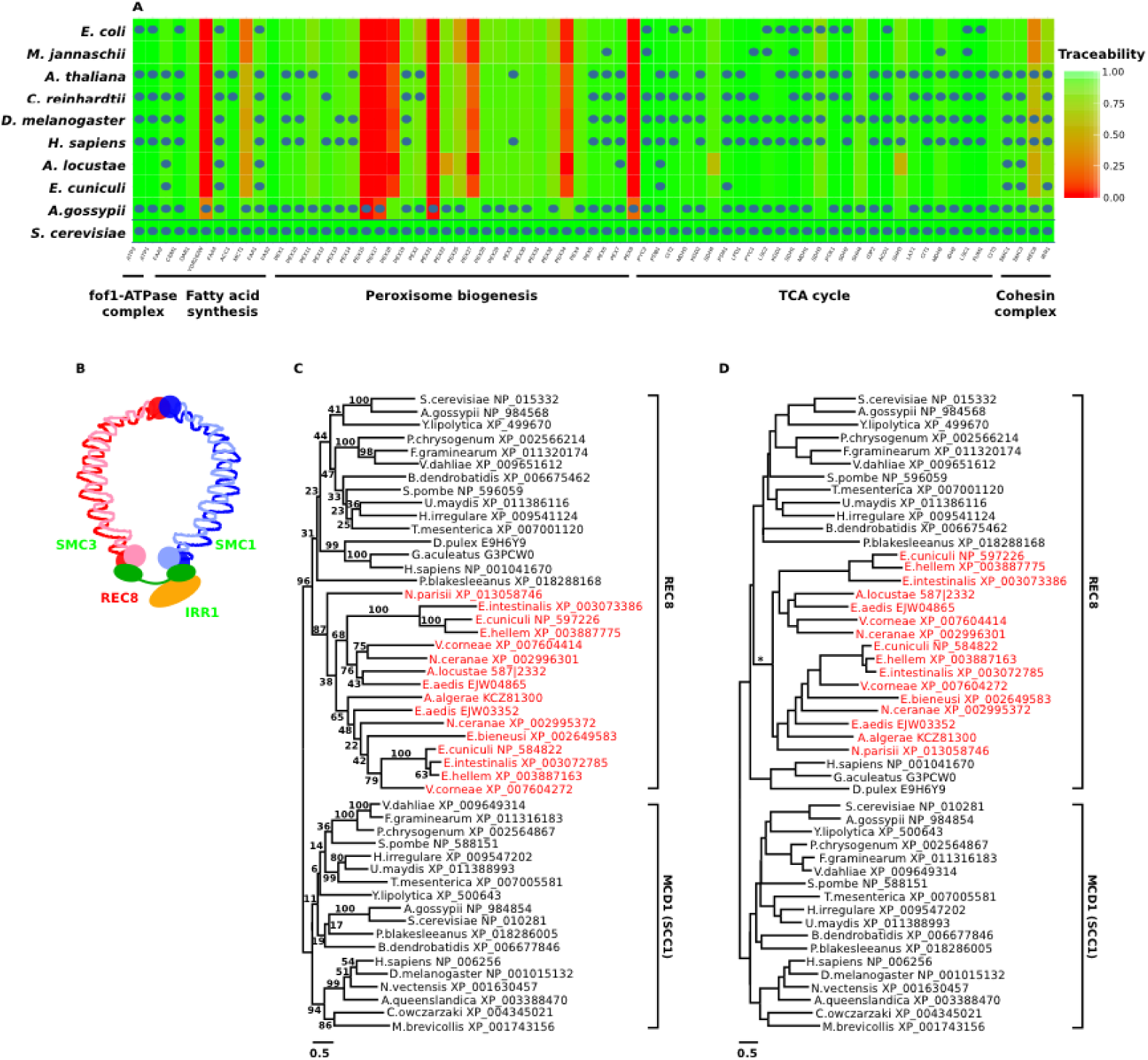
**A**, Phylogenetic profiles for the components of fungal key metabolic pathways across 10 representative species from the tree of life. The background colour gives the information of protein traceability ranging from green (high traceability) to red (low traceability). **B**, The four proteins of the yeast cohesin complex form a ring like structure. Font colour of the protein names indicate that the protein traceability in the microsporidium *E. cuniculi* is above (green) or below (red) 0.75. **C**, maximum likelihood tree of REC8 and MCD1 (syn. SCC1) orthologs. The microsporidian REC8 candidates are coloured in red. Branch labels represent percent bootstrap support. **D**, Alternative phylogeny for the REC8 / MCD1 (SCC1) protein family. It features monophyletic fungal REC8 and MCD1 (SCC1), respectively. The animal REC8 proteins are placed as sister to monophyletic fungal and microsporidian REC8 proteins. The branching orders in the fungal subtrees follows the accepted species phylogeny. The alternative tree is with a Δ_LogLikelihood_ = 25.7 not significantly worse than the ML tree shown in C (SH test: p>0.05). The asterisk indicates a gene duplication on the microsporidian linage that gave rise to the two paralogous microsporidian REC8 lineages.

The situation is different for proteins involved in meiosis and recombination. Yeast, as well as most other eukaryotes, share a conserved set of 29 proteins involved in this process (Malik, et al. 2007). Microsporidia, lack orthologs to several of these (Cuomo, et al. 2012) (*see* supplementary table S6). However, for three out of six absent orthologs the traceability of the yeast protein in microsporidia is low. This provides a clear indication that orthologs might have been overlooked. One protein, REC8, exemplifies the problem best. In yeast, REC8 forms with IRR1, SMC1, and SMC3 the cohesin complex, a ring-like structure that keeps sister chromatids connected during meiosis (Klein, et al. 1999) (fig. 6B). Interestingly, *E. cuniculi* harbours orthologs to three of the four genes (fig. 6A and *see* supplementary table S6). This raises the question about the whereabouts of the fourth member of this complex, REC8, which closes the ring-like structure. So far, a single report claims the presence of REC8 in the microsporidium *E. cuniculi* (Malik, et al. 2007). However, the search strategy that was used – a unidirectional Psi-Blast search (Altschul, et al. 1997) – lacks the precision to support this conclusion (Chen, et al. 2007). Consequently, a study based on ortholog searches reported the absence of this protein in *E. cuniculi*, and it identified *N. parisii* as the only microsporidian species harbouring an ortholog to the fungal REC8 (Cuomo, et al. 2012). To explain the sporadic presence of REC8 among microsporidia, Cuomo et al. (2012) hypothesized that the shorter period of time that *N. parisii* has been passaged in a laboratory setting, compared with other microsporidian species, caused the retention of REC8 only in this species. To resolve the controversy, we consulted the traceability of REC8 (fig. 6A). With a value of 0.5 in *E. cuniculi* it is substantially below the empirically determined critical value of 0.75. We took this as a reason for increasing the search sensitivity to identify highly diverged microsporidian REC8 orthologs, taking however the risk to end up with false positive predictions. In the first step, we screened the protein sets of 10 microsporidian species for sequences harbouring the Rad21_Rec8_N Pfam domain (PF04824), which occurs in REC8. This identified in six of the eleven species two proteins each, among them *E. cuniculi.* In each of the remaining four species only a single protein carried the PF04824 domain, among them *N. parisii.* We then extended the search to other eukaryotes (*see* supplementary fig. S10). Fungi, in general, possess two proteins with the PF04824 domain. In yeast, these correspond to REC8 and MCD1 (synonym SCC1). MCD1 is the protein that replaces REC8 in the cohesin complex during mitosis (Klein, et al. 1999). Thus, the identification of two microsporidian proteins with the Rad21_Rec8_N domains resembles the situation generally seen in fungi. However, at this step of the analysis, the precise identity of the microsporidian proteins remains unclear.

In the next step, we reconstructed the evolutionary relationships of a subset of fungal and non-fungal REC8 and MCD1 (SCC1) orthologs together with the microsporidian candidates (fig. 6C). Although this tree is not well resolved and renders, for example, the fungal REC8 proteins paraphyletic, it already supports a grouping of the microsporidian sequences with fungal and animal REC8 orthologs. Subsequently, we rearranged the tree topology to reflect the accepted evolutionary relationships of fungi, microsporidia and animals. A topology test revealed that the likelihood of the rearranged tree is with a Δ_LogLikelihood_ = 25.7 not significantly worse than the maximum likelihood tree (SH test: p>0.05 Shimodaira and Hasegawa (1999)). The data is therefore compatible with the hypothesis that microsporidian REC8 candidates form the sister clade of the fungal REC8 proteins, to the exclusion of the animal REC8 proteins (fig. 6D). Paired with the observation that the domain architecture of the microsporidian proteins agrees with that of yeast REC8 (*see* supplementary fig. S11), this indicates that we have indeed identified the missing REC8 orthologs in microsporidia.

In summary, the REC8 example shows that missing orthologs in the quickly evolving microsporidia are not always an effect of the rampant gene loss that is characteristic for this taxonomic group (Corradi and Slamovits 2011). Here, we provide for the first time convincing evidence that REC8 orthologs are widespread among microsporidia. The meiotic cohesin complex might therefore function in microsporidia as described for yeast. It should be noted, however, that we find no trace of MCD1 (SCC1), the mitotic counterpart of REC8. As this protein has a high traceability in the microsporidia, we propose a genuine gene loss of the Mcd1 gene (Supplementary Table 6). In this context, it is intriguing that we observe two paralogous REC8 proteins in the microsporidia, whose emergence via a gene duplication can be dated to the last common ancestor of the microsporidia. Notably, six out of ten microsporidian species harbour both paralogs. It is tempting to speculate that the apparent loss of the Mcd1 (Scc1) gene on the microsporidian lineage was compensated by a duplication of Rec8.

## Conclusion

Orthologs form the essential basis to propagate functional annotations between proteins of different species and to reconstruct the evolutionary past. So far, it has largely remained a matter of speculation to what extent limitations in the sensitivity of ortholog searches have influenced insights gained from these reconstructions. Here, we have presented a software, protTrace, facilitating a simulation-based procedure to assess the evolutionary traceability of a seed protein over time when using standard ortholog searches. In contrast to existing approaches, protTrace can infer constraints on the evolutionary sequence change of the seed protein from the presence of Pfam domains. This has two main advantages: The constraint estimates are independent from the availability and the phylogenetic diversity of orthologs to the seed protein; and the constraint pattern for a protein depends only on its Pfam domain composition and not on the species it was derived from. On the example of MRS2 we have shown that relying on Pfam for inferring the evolutionary constraints bears also a certain risk. Those sequences harbouring evolutionarily conserved regions that are not (yet) represented by a Pfam model will assume to evolve free of constraint. However, as Pfam becomes more comprehensive, the problem will ameliorate. Moreover, protTrace facilitates the inference of custom constraint patterns from a user-provided alignment of orthologs to the seed protein. The generally high traceability of enzymes indicates that orthologs are readily identifiable throughout the tree of life, explaining why ancestral gene set reconstructions is enriched for catalytic functions. This is contrasted by proteins preferably involved in regulatory processes, for which traceability implies that most signals informing about any ancient evolutionary origin have long been lost. Future attempts to unravel the remaining traces will have now the possibility to adapt the sensitivities of ortholog searches according to the traceabilities of the individual proteins. If the traceability of a protein is high, an increase of the search sensitivity – which naturally comes at the cost of a reduced specificity – is bound to result in false positive predictions. However, if the traceability is low, more sensitive searches may detect remaining faint signals of an evolutionary relationship between proteins in two species. In these cases, a careful downstream analysis including domain architecture comparison, phylogenetic tree reconstruction, and screen for interacting partners is then required to validate candidates resulting from such a relaxed search. Exemplarily for yeast REC8, we demonstrated that a limited traceability is indeed an issue that compromises ortholog detection and can lead to wrong evolutionary conclusions. Contrary to current belief, we could show that REC8 is present and widespread in microsporidia, rendering the cohesin complex complete and probably functional. Thus, microsporidia bring along the necessary prerequisite for both meiosis and recombination.

In summary, the evolutionary traceability of proteins brings us one step closer towards deciding when the absence of evidence for an ortholog is evidence for its absence, and when it is not (Alderson 2004).

## Materials and Methods

### Data sets

Our analyses are based on 232 species representing the three domains of life (*see* supplementary table S1). The phylogenetic tree for these species was obtained from NCBI CommonTree (https://www.ncbi.nlm.nih.gov/Taxonomv/CommonTree/wwwcmt.cgi).

The Last Universal Common Ancestor (LUCA) gene sets (1203 genes) were downloaded from LUCApedia (Goldman, et al. 2013), a database consisting of all LUCA gene sets proposed by different studies. The essential genes set (1110 genes) for *S. cerevisiae* was obtained from Database of Essential Genes (DEG) (Luo, et al. 2014). The LUCA genes and the essential genes are listed in supplementary table S3.

### Compilation of orthologous groups

First, orthologs for the seed protein are retrieved from the corresponding ortholog group provided by the OMA database (Altenhoff, et al. 2015). We then extend the OMA ortholog group with sequences from a collection of 232 species (*see* supplementary table S1) using HaMStR (Ebersberger, et al. 2009), a profile hidden Markov model (pHMM) based ortholog search tool. HaMStR was run on with the following parameters: -*strict*, -*checkCoorthologsRef*, -*hit*_*limit*=*1* and -*representative*. For query proteins without orthologs in the OMA database, we directly perform a targeted ortholog search using HaMStR-OneSeq (Ebersberger, et al. 2014) in the gene sets of 232 species. HaMStR-OneSeq is an extended version of HaMStR that compiles in an iterative procedure an initial core-ortholog set for pHMM training. Once the training is completed, a final ortholog search in all taxa concludes the procedure. HaMStR-OneSeq is run with the following parameters: -*coreOrth*=*5*, -*minDist*=*genus*, -*maxDist*=*superkingdom*, -*checkCoorthologsRef*, -*strict* and -*rep.* Alternatively, we used ortholog groups provided by OrthoDB (Zdobnov, et al. 2017) for parameterizing the evolutionary models. Comparing the two alternative ways to compile the training data across the entire yeast proteome reveals highly correlated traceability values (Pearson’s *r*=0.92). This shows that the choice of the ortholog search algorithm has only little influence on the traceability estimates. Likewise, traceability estimates remain largely unaffected when confining the training data for the evolutionary parameters to fungal orthologs rather than recruiting orthologs, where available, from the entire tree of life (see supplementary fig. S5).

### Maximum likelihood distance estimation

We computed pairwise maximum likelihood distances between proteins using TreePuzzle v5.225 (Schmidt, et al. 2002). To arrive at an average maximum likelihood genetic distance between any pair of species, we extracted and aligned all pairwise orthologs for the two species from the OMA database (Altenhoff, et al. 2015). In the case of 1 to many ortholog groups, we considered all induced pairwise orthology relationships. The alignments were then concatenated and served as input for TreePuzzle to compute an average maximum likelihood distance. The procedure was repeated for all species pairs in the reference tree to obtain an all-against-all maximum likelihood distance matrix.

### Annotation of Pfam domain

We annotated Pfam (Finn, et al. 2016) domains using *hmmscan* (Finn, et al. 2011) with parameters ‐‐*notextw* and -*E 0.01*.

### GO term enrichment analysis

We searched for GO terms enriched in a set of yeast proteins with GOrilla (Eden, et al. 2009). The entire gene set of *S. cerevisiae* served as the background set. An E-value cut-off of 10E-3 was applied. Significantly enriched GO terms were then visualized using Revigo (Supek, et al. 2011).

### Phylogenetic analysis of REC8

The domain annotation of REC8 in yeast (*S. cerevisiae*) revealed the presence of a Rad21_REC8_N domain (PF04825). Using the Rad21_REC8_N profile HMM obtained from Pfam (Finn, et al. 2016), we searched with *hmmsearch* (Finn, et al. 2011) for proteins harbouring this domain in the gene sets of ten microsporidia (*E. cuniculi*, *A. locustae*, *N. ceranae*, *E. bieneusi*, *E. aedis*, *E. hellem*, *E. intestinalis*, *A. algerae*, *V. corneae* and *N. parisii*) and of yeast. The search in yeast resulted in a second protein, MCD1/Scc1, also containing the Rad21_REC8_N domain. We then retrieved REC8 and MCD1/Scc1 orthologs from training data used for the traceability calculation in the following fungal and outgroup species - *A. gossypii*, *Y. lipolytica*, *F. graminearum*, *V. dahliae*, *P. chrysogenum*, *S. pombe*, *T. mesenterica*, *U. maydis*, *H. irregulare*, *P. blakesleeanus*, *B. dendrobaditis*, *C. owczarzaki*, *M. brevicollis*, *A. queenslandica*, *N. vectensis*, *D. melanogaster*, and *H. sapiens.* Because both OMA and HaMStR found no orthologs to yeast REC8 in animals, we complemented the data with the *H. sapiens* REC8 protein (NCBI accession: NP_001041670), and its InParanoid (Ostlund, et al. 2010) orthologs from *G. aculeatus*, and *D. pulex.* All sequences were aligned with MAFFT v7.304 using the option *L*-*INS*-*i*. From the resulting multiple sequence alignment, we computed a maximum likelihood (ML) tree with 100 bootstraps run using RAxML v8 (Stamatakis 2014), modelling the substitution process with *PROTGAMMALG*, the best model obtained from ProtTest v3 (Abascal, et al. 2005). Tree topology testing was performed using the routines implemented in RAxML. Pfam domain architecture display on a phylogenetic tree was done with doMosaics (Moore, et al. 2014).

## Data availability

All data that supports the finding of this study are available from the corresponding author upon request.

## Supplementary Information

is available in the online version of the paper.

## Acknowledgements

We thank Tina Koestler for her contributions in the early stage of the project, and many colleagues, among them Kathi Zarnack and Michael Hofreiter, for their comments on the manuscript. This work was supported by the Marie Curie ITN project CALIPSO (GA ITN-2013 607 607), by the Deutsche Forschungsgemeinschaft (DFG FOR 2251; Project Grant EB 285/2-1), and by the Austrian Science Fund (FWF Grant to A.v.H. I 1824-B22). A.v.H. also acknowledges support from the Medical University of Vienna and the University of Vienna.

